# Killing from the inside: Intracellular role of T3SS in the fate of *Pseudomonas aeruginosa* within macrophages revealed by *mgtC* and *oprF* mutants

**DOI:** 10.1101/389718

**Authors:** Preeti Garai, Laurence Berry, Malika Moussouni, Sophie Bleves, Anne-Béatrice Blanc-Potard

## Abstract

While considered solely an extracellular pathogen, increasing evidence indicates that *Pseudomonas aeruginosa* encounters intracellular environment in diverse mammalian cell types, including macrophages. In the present study, we have deciphered the intramacrophage fate of wild-type *P. aeruginosa* PAO1 strain by live and electron microscopy. *P. aeruginosa* first resided in phagosomal vacuoles and subsequently could be detected in the cytoplasm, indicating phagosomal escape of the pathogen, a finding also supported by vacuolar rupture assay. The intracellular bacteria could eventually induce cell lysis. Two bacterial factors, MgtC and OprF, recently identified to be important for survival of *P. aeruginosa* in macrophages, were found to be involved in bacterial escape from the phagosome as well as cell lysis caused by intracellular bacteria. Strikingly, type III secretion system (T3SS) genes of *P. aeruginosa* were down-regulated within macrophages in both *mgtC* and *oprF* mutants. Concordantly, cyclic di-GMP (c-di-GMP) level was increased in both mutants, providing a clue for negative regulation of T3SS inside macrophages. Consistent with the phenotypes and gene expression pattern of *mgtC* and *oprF* mutants, a T3SS mutant *(ΔpscN)* exhibited defect in phagosomal escape and macrophage lysis driven by internalized bacteria. Importantly, these effects appeared to be largely dependent on the ExoS effector, in contrast with the known T3SS-dependent, but ExoS independent, cytotoxicity caused by extracellular *P. aeruginosa* towards macrophages. Hence, our work highlights T3SS and ExoS, whose expression is modulated by MgtC and OprF, as key players in the intramacrophage life of *P. aeruginosa*, allowing internalized bacteria to evade macrophages.

**Author summary:** The ability of professional phagocytes to ingest and kill microorganisms is central to host defense and *Pseudomonas aeruginosa* has developed mechanisms to avoid being killed by phagocytes. While considered an extracellular pathogen, *P. aeruginosa* has been reported to be engulfed by macrophages in animal models. Here, we visualized the fate of *P. aeruginosa* within cultured macrophages, revealing macrophage lysis driven by intracellular *P. aeruginosa*. Two bacterial factors, MgtC and OprF, recently discovered to be involved in the intramacrophage survival of *P. aeruginosa*, appeared to play role in this cytotoxicity caused by intracellular bacteria. We provided evidence that type III secretion system (T3SS) gene expression is lowered intracellularly in *mgtC* and *oprF* mutants. We further showed that intramacrophage *P. aeruginosa* uses its T3SS, specifically the ExoS effector, to promote phagosomal escape and cell lysis. We thus describe a transient intramacrophage stage of *P. aeruginosa* that could contribute to bacterial dissemination.

## Introduction

Pathogenic bacteria are commonly classified as intracellular or extracellular pathogens [1]. Intracellular bacterial pathogens, such as *Mycobacterium tuberculosis* or *Salmonella enterica*, can replicate within host cells, including macrophages. In contrast, extracellular pathogens, such as *Staphylococcus aureus*, *Pseudomonas aeruginosa* or streptococci, avoid phagocytosis or exhibit cytotoxicity towards phagocytic cells, thus promoting extracellular multiplication. However, recent data have emphasized the fact that several extracellular pathogens can enter host cells *in vivo*, resulting in a phase of intracellular residence, which can be of importance in addition to the typical extracellular infection. For example, once considered an extracellular pathogen, it is now established that *S. aureus* can survive within many mammalian cell types including macrophages [2,3] and the intramacrophage fate of *S. aureus* has been deciphered [4,5]. Moreover, an intracellular phase within splenic macrophages has been recently shown to play a crucial role to initiate dissemination of *Streptococcus pneumonia*, providing a divergence from the dogma that considered this bacterial pathogen as the classical example of extracellular pathogen [6].

The environmental bacterium and opportunistic human pathogen *P. aeruginosa* is responsible for a variety of acute infections and is a major cause of mortality in chronically infected cystic fibrosis (CF) patients. The chronic infection of *P. aeruginosa* and its resistance to treatment is largely due to its ability to form biofilms, which relies on the production of exopolysaccharides (EPS), whereas the type III secretion system (T3SS) is reported to play a key role in the pathogenesis of acute *P. aeruginosa* infections [7]. Four T3SS effectors (ExoU, ExoS, ExoT, ExoY) have been identified, ExoS being more prevalent and nearly always mutually exclusive with the potent cytotoxin ExoU [8–10]. An intracellular step in airway epithelial cells has been proposed to occur before the formation of biofilm during the acute phase of infection [11,12] and intracellular stage of *P. aeruginosa* within cultured epithelial cells has been fairly studied [13–15]. The principle that *P. aeruginosa* is not exclusively an extracellular pathogen has been convincingly established by the recent use of advanced imaging methods to track bacteria within epithelial cells [16]. The T3SS, more specifically its effector ExoS, has been implicated in the formation of membrane blebs-niche and avoidance of acidified compartments to allow bacterial multiplication within epithelial cells [16–18]. Concordantly, T3SS was recently shown to be expressed in *P. aeruginosa* internalized in epithelial cells [16]. Regarding the interaction with macrophages, *P. aeruginosa* has developed mechanisms to avoid phagocytosis [19]. However, *P. aeruginosa* has been shown to be phagocytosed by macrophages in an acute model of infection in zebrafish embryos [20] and has been reported to be engulfed by alveolar macrophages in the lungs of mice early after pulmonary infection [21], suggesting that *P. aeruginosa* may have to escape macrophage killing. Although *P. aeruginosa* has been localized within cultured macrophages during gentamicin protective assays [22–24], the intramacrophage fate of the bacteria has not been characterized and bacterial factors involved in this step remain largely unexplored.

Bacterial survival in a particular niche requires the development of an adaptive response, generally mediated by regulation of the genes involved in physiological adaptation to the microenvironment. The identification of mutants lacking the ability to survive within macrophages and the study of *P. aeruginosa* gene expression inside macrophages is critical to determine bacterial players in this step. We have previously uncovered MgtC as a bacterial factor involved in the intramacrophage survival of *P. aeruginosa* [23,25]. In agreement with this intramacrophage role, expression of *Pseudomonas mgtC* (PA4635) gene is induced when the bacteria reside inside macrophages [23]. MgtC is known to promote intramacrophage growth in several classical intracellular bacteria, including *Salmonella typhimurium* where it inhibits bacterial ATP synthase and represses cellulose production [26–28], and is considered as a clue to reveal bacterial pathogens adapted to an intramacrophage stage [28,29]. In addition, a recent study has implicated OprF in the ability of otopathogenic *P. aeruginosa* strains to survive inside macrophages [24]. OprF is an outer membrane porin that can modulate the production of various virulence factors of *P. aeruginosa* [30,31].

In the present study, we investigated the fate of *P. aeruginosa* within macrophages using wild-type PAO1 strain, which lacks ExoU, as well as *mgtC* and *oprF* mutants. We also explored, for the first time, the regulation of T3SS genes when *P. aeruginosa* resides inside macrophages. The T3SS and ExoS effector, whose expression was found to be modulated by MgtC and OprF intracellularly, were seemingly involved in this intracellular stage, playing a role in vacuolar escape and cell lysis caused by intracellular bacteria.

## Results

### Intracellular *P. aeruginosa* can promote macrophage lysis

We previously visualized fluorescent *P. aeruginosa* residing within fixed macrophages [23]. To investigate the fate of *P. aeruginosa* after phagocytosis in a dynamic way, we monitored macrophages infected with fluorescent bacteria using time-lapse live microscopy. J774 macrophages were infected with wild-type PAO1 strain expressing GFP (Multiplicity of infection or MOI=10). After 25 minutes of phagocytosis, several washes were performed to remove adherent bacteria and gentamicin was added to kill extracellular bacteria. Microscopic observation of infected macrophages up to 3 hrs showed cell lysis with time (Fig 1), which can be attributed to intracellular bacteria since gentamicin was retained throughout the experiment. No lysis of uninfected cells present in the same field was observed (Fig 1 and S1 Fig), as expected from an event due to intracellular bacteria rather than extracellular bacteria. The cell lysis appeared to be a rapid event as shown by the movie (S1 Movie).

**Fig 1.**
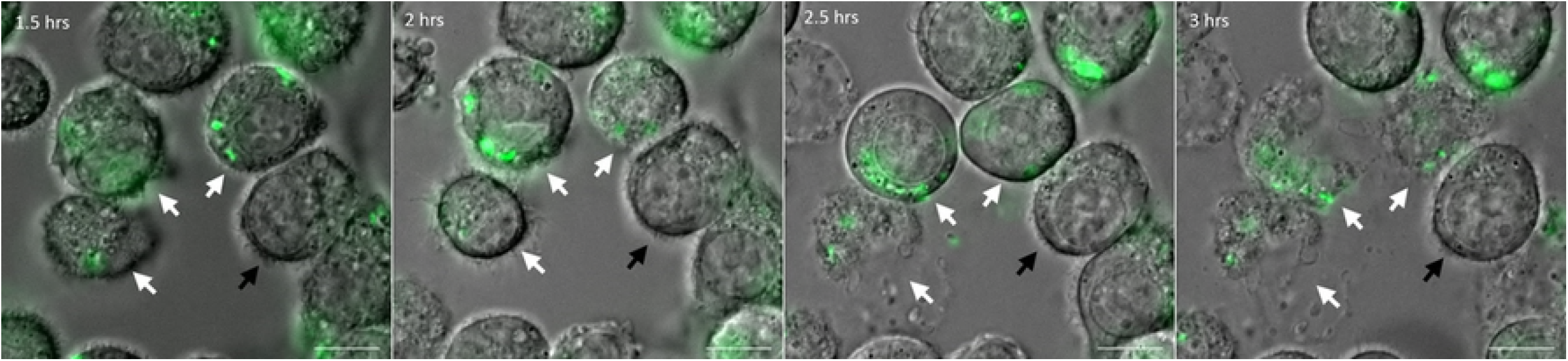
Live imaging of macrophages infected with *P. aeruginosa.* J774 macrophages were infected with PAO1 wild-type (WT) strain expressing GFP. Time lapse imaging was started at 1.5 hrs post-phagocytosis. Cells were maintained in DMEM supplemented with gentamicin at 37°C and 5% CO_2_ throughout imaging. White arrows point at the cells that harbor intracellular bacteria and undergo lysis between 1.5 hrs and 3 hrs post-phagocytosis. Black arrow shows an uninfected and unlysed cell. Scale bar is equivalent to 10 μm.

We further examined intracellular *P. aeruginosa* within macrophages in more detail using transmission electron microscopy (TEM). J774 macrophages infected with wild-type PAO1 strain, were subjected to fixation at early time post-infection (30 min after phagocytosis) or at later time of infection (3 hrs after phagocytosis). After phagocytosis, *P. aeruginosa* was present in membrane bound vacuoles inside macrophages (Fig 2A). At a later time point, some bacteria could be found in the cytoplasm with no surrounding membrane, suggesting disruption of the vacuole membrane (Fig 2B). The infected macrophage was damaged, displaying no pseudopodia, highly condensed chromatin and membrane blebbing. Healthy infected cells were also observed, where bacteria were mostly found in vacuoles partially or totally filled with heterogeneous electron dense material, suggesting that the vacuole has fused with lysosomes (Fig 2C). We further examined the association between fluorescent PAO1 bacteria and the LysoTracker probe during infection using fixed macrophages. Bacteria colocalizing with LysoTracker could be visualized (S2 Fig), thereby corroborating the TEM observation of vacuoles that have presumably fused with lysosomes with the localization of bacteria in acidified compartment.

**Fig 2.**
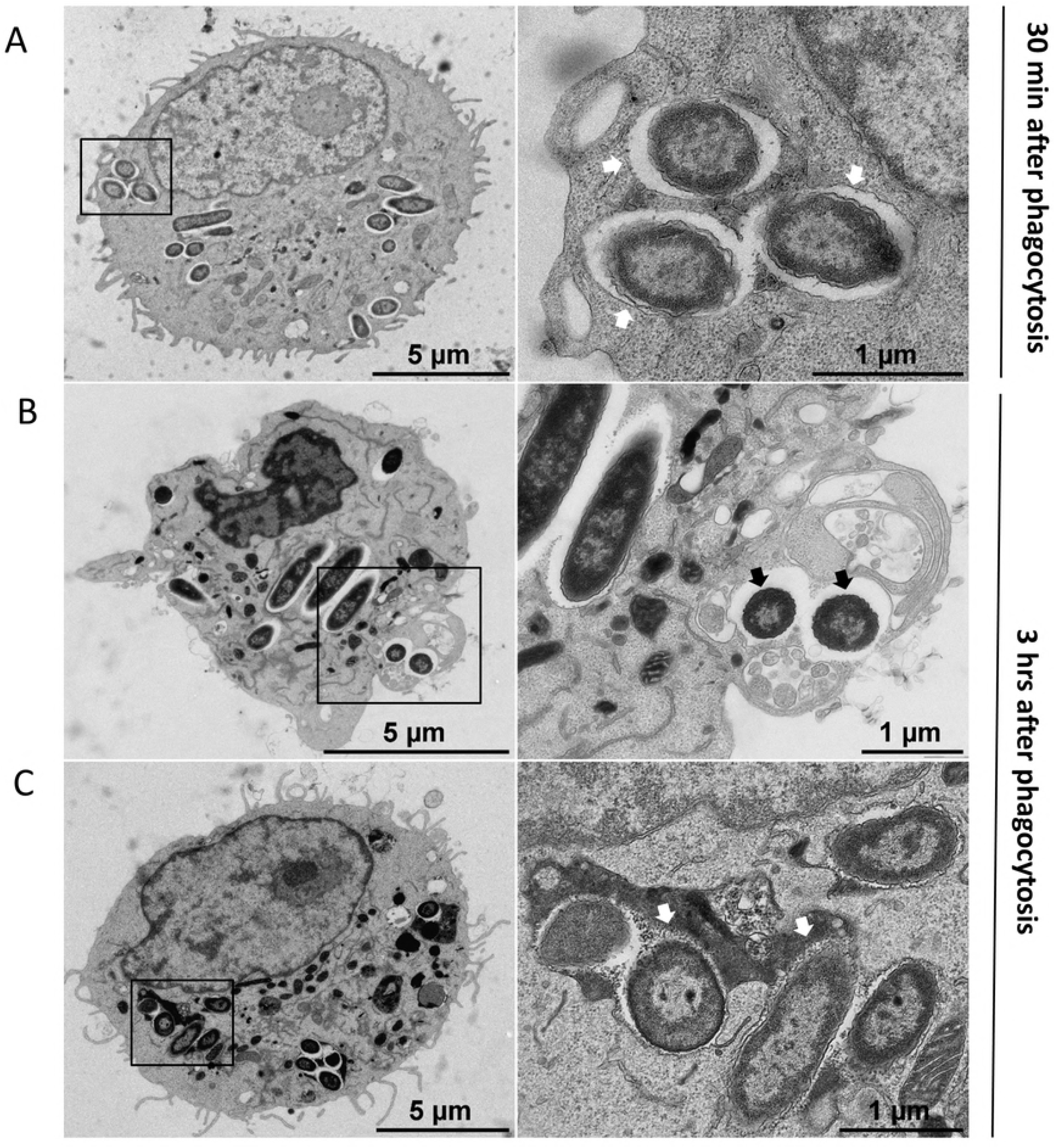
Transmission electron micrographs (TEM) of *P. aeruginosa* within macrophages. J774 macrophages were infected with *P. aeruginosa* for 30 min (A) or 3 hrs (B and C) and subjected to TEM (left panels). Black rectangles show intracellular bacteria that are shown at higher magnification in the right panels. At early time after phagocytosis (A), most of bacteria were found inside membrane bound vacuoles (white arrows). B. At later time, some bacteria can be observed in the cytoplasm with no surrounding membrane suggesting disruption of the vacuole membrane (black arrows). The infected macrophage in panel B shows an abnormal morphology, with no pseudopodia, highly condensed chromatin, and membrane blebbing. C. At later time, bacteria can also be found in vacuole partially or totally filled with heterogeneous electron dense material (white arrows), suggesting that the vacuole has fused with lysosomes. The infected macrophage shown in panel C appears normal.

TEM analyses allowed us to conclude that wild-type PAO1 strain has the ability to reside within vacuoles and possibly escape from the phagosome into the cytoplasm and promote cell damage. A rapid cell lysis event caused by intracellular *P. aeruginosa* visualized by live microscopy revealed that phagocytosed bacteria can escape from macrophage through cell lysis.

### Intracellular *mgtC* and *oprF* mutants are compromised in cell lysis

Our previous results based on gentamicin protection assay on J774 infected macrophages and counting of colony forming units (CFU) indicated that *mgtC* mutant in the PAO1 background survived to a lesser extent than the wild-type strain [23]. More recently, an *oprF* mutant in a *P. aeruginosa* otopathogenic strain was also found to be defective in intracellular survival in mouse bone marrow macrophages based on gentamicin protection assay [24]. To analyze phenotypes of *oprF* mutant towards J774 macrophages and compare with *mgtC* mutant, we used here a previously described *oprF* mutant in the PAO1 parental background [23,32]. Similarly to *mgtC* mutant, PAO1 *oprF* mutant was found deficient for intracellular survival in J774 macrophages 2 hrs after phagocytosis comparatively to wild-type PAO1 (S3 Fig).

Based on the finding that intracellular *P. aeruginosa* can cause macrophage lysis, we developed a suitable assay using fluorescent microscopic analysis to quantify cell damage caused by intracellular bacteria. Macrophages were infected with fluorescent bacteria, treated with gentamicin for 2 hrs after phagocytosis and fixed cells were stained with phalloidin-TRITC to label F-actin and visualize macrophage morphology. A clear cortical red labeling was seen in most cells, which is indicative of the plasma membrane-associated cortical actin. Few infected cells appeared lysed due to the loss of cortical actin staining (Fig 3A), which agrees with the observation of lysed macrophages in time lapse fluorescence microscopy (Fig 1). The loss of cortical actin staining was due to internalized bacteria because upon arresting phagocytosis, by treating macrophages with cytochalasin D, bacteria were not internalized and loss of cortical actin staining was not observed (S4 Fig). Quantification of the number of cells without phalloidin labelling showed highest value for wild-type strain, intermediate for *mgtC* mutant and lowest for *oprF* mutant (Fig 3B). Thus, both intracellular *mgtC* and *oprF* mutants appear compromised for cell lysis. This feature is not linked to a lower internalization rate of the mutant strains, since *mgtC* mutant is more phagocytozed than wild-type strain [23] and a similar trend was found for *oprF* mutant (not shown). Since T3SS has been involved in the intracellular life of *P. aeruginosa* in other cell types [16], we next monitored the expression of T3SS genes during the residence of bacteria within macrophages.

**Fig 3.**
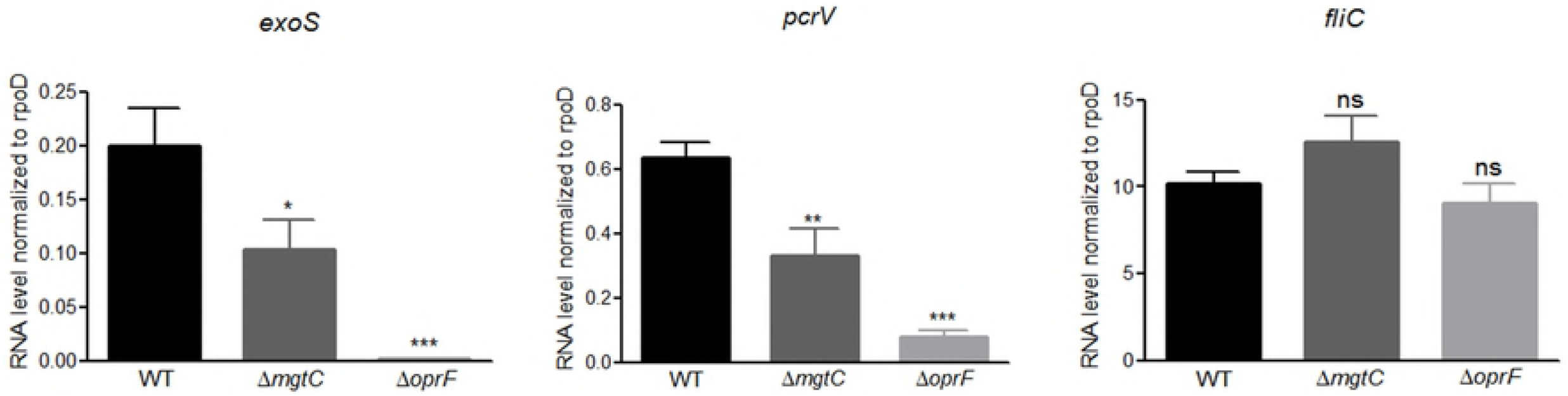
Visualization (A) and quantification (B) of lysed infected cells upon phalloidin labeling. GFP expressing PAO1 WT, *ΔmgtC* and *ΔoprF* strains were used for infecting J774 macrophages. Gentamicin was added after phagocytosis and cells were fixed at 2 hrs postphagocytosis, stained with phalloidin and imaged with fluorescent microscope. DAPI was used to stain the nucleus. Cells that have intracellular bacteria, but lack the phalloidin cortical label were considered as lysed by intracellular bacteria (shown by arrows). Scale bar is equivalent to 10 μm. After imaging, infected cells were counted and percentage of lysed cells with intracellular bacteria out of total number of cells was plotted for each strain. Error bars correspond to standard errors from three independent experiments. At least 200 cells were counted per strain. The asterisks indicate *P* values (One way ANOVA, where all strains were compared to WT using Dunnet’s multiple comparison post-test, \**P* <0.05 and \*\*\**P* <0.001), showing statistical significance with respect to WT.

### Down-regulation of T3SS genes expression in *mgtC* and *oprF* mutants inside macrophages

The *P. aeruginosa oprF* mutant was reported to be defective in the secretion of T3SS effectors in liquid culture and in the production of PcrV, the T3SS needle tip protein [30,32], but the effect of OprF on transcription of T3SS genes has not been tested so far. We thus investigated the expression of the effector gene *exoS* and of *pcrV* in *oprF* mutant strain in comparison to wild-type strain upon macrophage infection. As a control, we tested *fliC,* which is not part of the T3SS regulon, although FliC was proposed to be secreted by the T3SS [33]. Expression of *pcrV* and *exoS* genes was highly reduced in the *oprF* mutant (Fig 4). Strikingly, the *mgtC* mutant also exhibited significantly reduced expression of these two T3SS genes (Fig 4), although to a lesser extent than the *oprF* mutant, indicating an unexpected interplay between MgtC and T3SS. On the other hand, *fliC* expression was not altered in both mutants.

**Fig 4.**
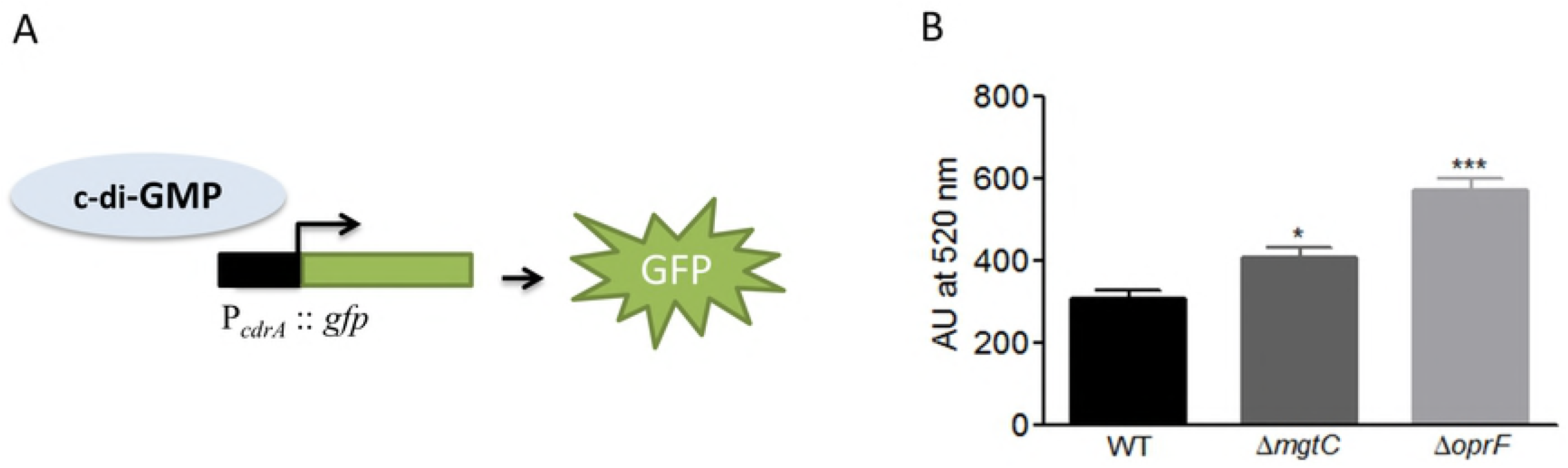
Expression of T3SS genes in *P. aeruginosa* strains residing in J774 macrophages. J774 macrophages were infected with PAO1 WT, *ΔmgtC* and *ΔoprF* strains. After phagocytosis, cells were maintained in DMEM supplemented with gentamicin. RNA was extracted from bacteria isolated from infected macrophages 1 hr after phagocytosis. The level of *exoS, pcrV* and *fliC* transcripts relative to those of the house-keeping gene *rpoD* was measured by qRT-PCR and plotted on the Y-axis. Error bars correspond to standard errors from at least three independent experiments. The asterisks indicate *P* values (One way ANOVA, where all strains were compared to WT using Dunnet’s multiple comparison posttest, *P <0.05, **P <0.01, ***P <0.001 and ns = *P* >0.05 or non-significant), showing statistical significance with respect to WT.

To address the mechanism behind the downregulation of transcription of T3SS in the *oprF* and *mgtC* mutant strains, we evaluated the level of the second messenger c-di-GMP, as it is known to participate in T3SS repression in *P. aeruginosa* [34]. The *oprF* mutant was already reported to have high production of c-di-GMP in liquid culture [32]. We used a fluorescence-based reporter that has been validated to gauge c-di-GMP level inside *P. aeruginosa* [35]. The pCdrA::*gfp* plasmid was introduced into wild-type and mutant strains and fluorescence was measured in infected macrophages (Fig 5). Both *oprF* and *mgtC* mutants exhibited significant increased activity of the *cdrA* promoter comparatively to wild-type strain, indicative of higher levels of c-di-GMP than the wild-type strain. Taken together, these results indicate that the production of c-di-GMP is increased relatively to wild-type strain in both *oprF* and *mgtC* mutants when bacteria reside within macrophages, thus providing a mechanistic clue for the negative effect on T3SS gene expression.

**Fig 5.**
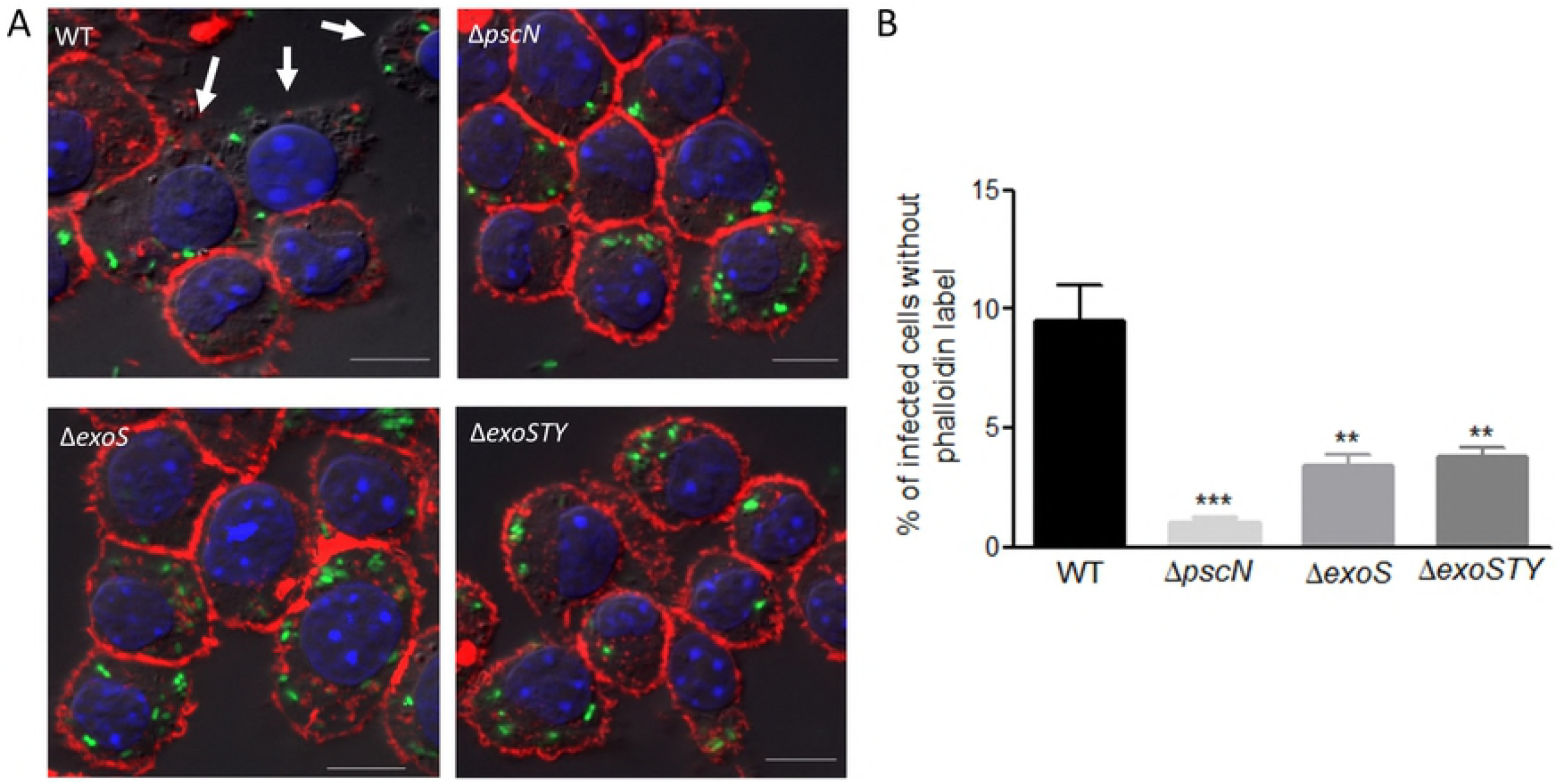
Measurement of c-di-GMP level in *ΔmgtC* and *ΔoprF* mutants inside macrophages. PAO1 WT, *ΔmgtC* and *ΔoprF* harboring reporter plasmid pCdrA::*gfp*, which expresses GFP under the control of the promoter of c-di-GMP responsive gene *cdrA* (A), were used to infect J774 cells. After phagocytosis, DMEM containing 300 μg/ml of amikacin was added to eliminate extracellular bacteria. Fluorescence (excitation, 485 nm and emission, 520 nm) was measured 1 hour after phagocytosis and plotted as arbitrary units (AU). Error bars correspond to standard errors from four independent experiments. The asterisks indicate *P* values (One way ANOVA, where all strains were compared to WT using Dunnet’s multiple comparison post-test, \**P* <0.05 and \*\*\**P* <0.001), showing statistical significance with respect to WT (B).

### A T3SS mutant is defective for cell death driven intracellularly in an ExoS-dependent manner

Given the effect of both *oprF* and *mgtC* deletions on the expression of T3SS genes within macrophages, we investigated the fate of a T3SS mutant upon phagocytosis. We first used a *pscN* mutant that is defective for the ATPase of the T3SS machinery [36]. Intracellular T3SS mutant did not induce loss of cortical phalloidin staining, as shown by fluorescent imaging and subsequent quantification, indicating lack of cell lysis (Fig 6A and B). To address the implication of T3SS effector proteins more specifically, we used *exoS* and *exoSTY* mutants. The intracellular lysis of macrophages was found to be reduced for the triple mutant *exoSTY* and to a similar extent for the *exoS* mutant (Fig 6A and B). Although the lysis by *exoS* mutant strain was substantially higher than that of *pscN* mutant, these results suggest that the T3SS-mediated intracellular lysis of macrophages by *P. aeruginosa* relies mainly on the T3SS effector ExoS. This cytotoxic effect mediated by intracellular bacteria thus differs from the classical T3SS-dependent cytotoxity driven by extracellular bacteria towards macrophages that has been reported to be mostly independent of ExoS [37–39]. To confirm this difference between intracellular and extracellular bacteria mediated lysis, we measured the LDH release upon an infection carried out without removing extracellular bacteria. As expected, similar values were obtained for wild-type strain and *exoS* mutant, whereas a *pscN* mutant showed significantly reduced LDH release (S5 Fig), which agrees with the reported T3SS-dependent but ExoS independent cytotoxicity caused by extracellular bacteria. Hence, our results indicate that, in contrast to extracellular *P. aeruginosa*, intracellular *P. aeruginosa* uses an ExoS-dependent T3SS mediated mechanism to promote macrophage lysis. Moreover, the phenotypes of T3SS mutants are consistent with that of *mgtC* and *oprF* mutants, in correlation with their level of T3SS gene expression.

**Fig 6.**
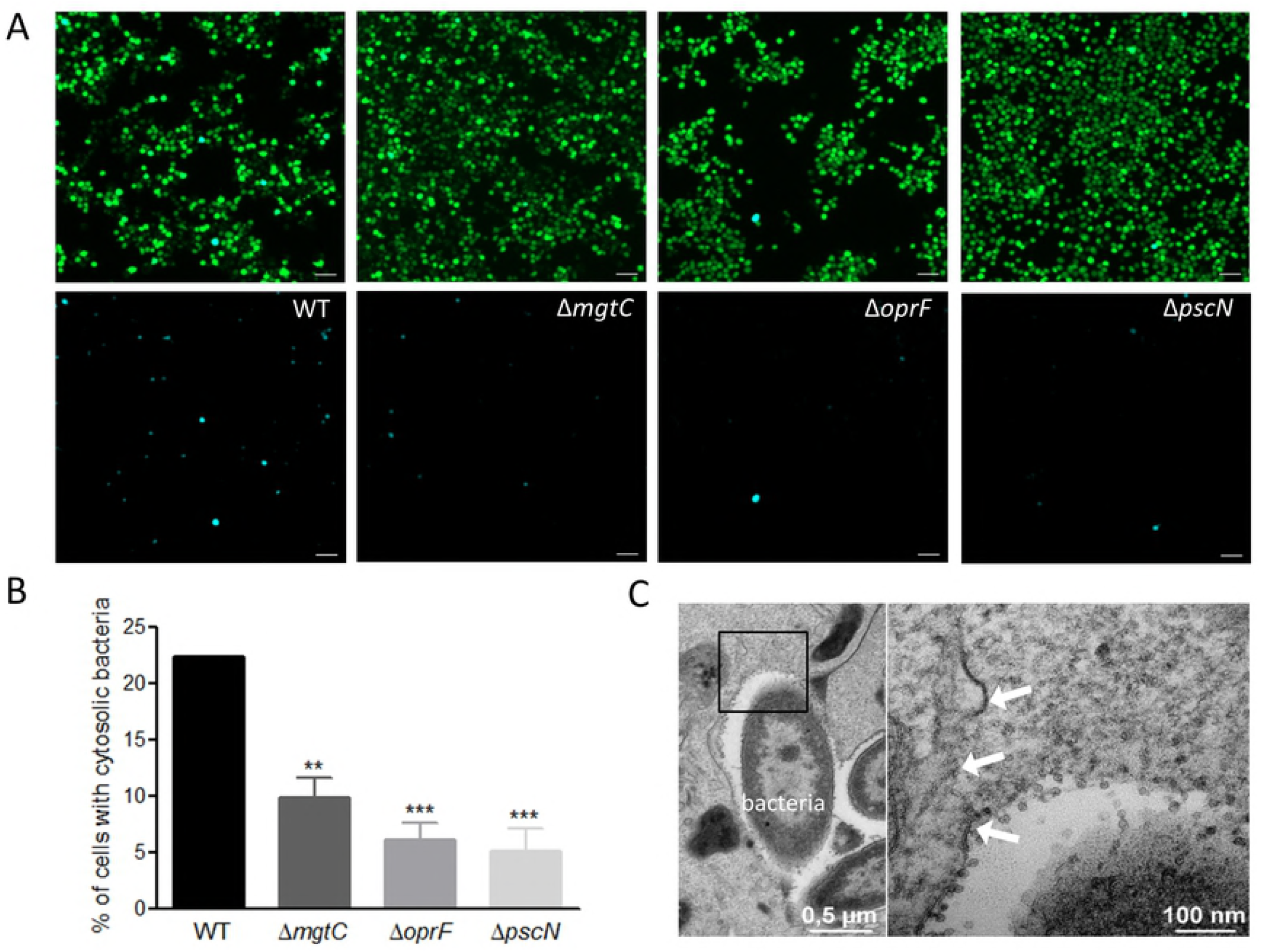
Role of T3SS in cell lysis induced by intracellular bacteria. J774 macrophages were infected with PAO1 WT, *ΔpscN, ΔexoS* and *ΔexoSTY* strains. After phagocytosis, cells were maintained in DMEM supplemented with gentamicin. Cells were imaged 2 hrs postphagocytosis after staining with phalloidin (A) and lysis was quantified (B) by counting infected cells lacking cortical labeling (indicated by arrows). Scale bar is equivalent to 10 μm. Percentage of lysed cells with intracellular bacteria out of total number of cells was plotted. Error bars correspond to standard errors from at least three independent experiments. At least 200 cells were counted per strain. The asterisks indicate *P* values (One way ANOVA, where all strains were compared to WT using Dunnet’s multiple comparison test, \*\**P* <0.01 and \*\*\**P* <0.001), showing statistical significance with respect to WT.

### P. aeruginosa T3SS, *oprF* and *mgtC* mutants displayed lower vacuolar escape than wild-type PAO1

The phagosomal environment of the macrophage is hostile for bacterial pathogens and TEM analysis indicated that PAO1 can be found in the cytoplasm, suggesting escape from the phagosomal vacuole (Fig 2B). We decided to address the intracellular role of T3SS, OprF and MgtC in the phagosomal escape of *P. aeruginosa* to the cytoplasm of macrophages. To monitor *P. aeruginosa* escape from phagosome, we used the CCF4-AM/β-lactamase assay that has been developed for tracking vacuolar rupture by intracellular pathogens [40]. This assay takes advantage of the natural production of β-lactamase by *P. aeruginosa* [41], which can cleave a fluorescent β-lactamase substrate, CCF4-AM, that is trapped within the host cytoplasm. CCF4-AM emits a green fluorescence signal, whereas in the presence of β-lactamase activity, a blue fluorescence signal is produced. The presence of blue fluorescent cells at 2 hrs post-phagocytosis indicated bacterial escape from the phagosome to the cytoplasm (Fig 7A). The escape of wild-type strain was compared to that of *mgtC, oprF* and *pscN* mutants by quantifying the percentage of blue fluorescent cells out of total green fluorescent cells. Wild-type strain showed significantly higher percentage of phagosomal escape than all mutants tested (Fig 7B). In agreement with the finding of phagosomal escape by CCF4-AM/β-lactamase assay, ruptured phagosomal membrane could be visualized by TEM in macrophages infected with PAO1 wild-type strain (Fig 7C). Both *exoS* and *exoSTY* effector mutants also displayed a low percentage of phagosomal escape, which appeared similar to that of *pscN* mutant (S6 Fig). These results indicate that the T3SS, through the ExoS effector, plays role in the escape of *P. aeruginosa* from the phagosome to the cytoplasm. The effect of MgtC and OprF in this process may as well be mediated by their effect on T3SS gene expression (Fig 4).

**Fig 7.**
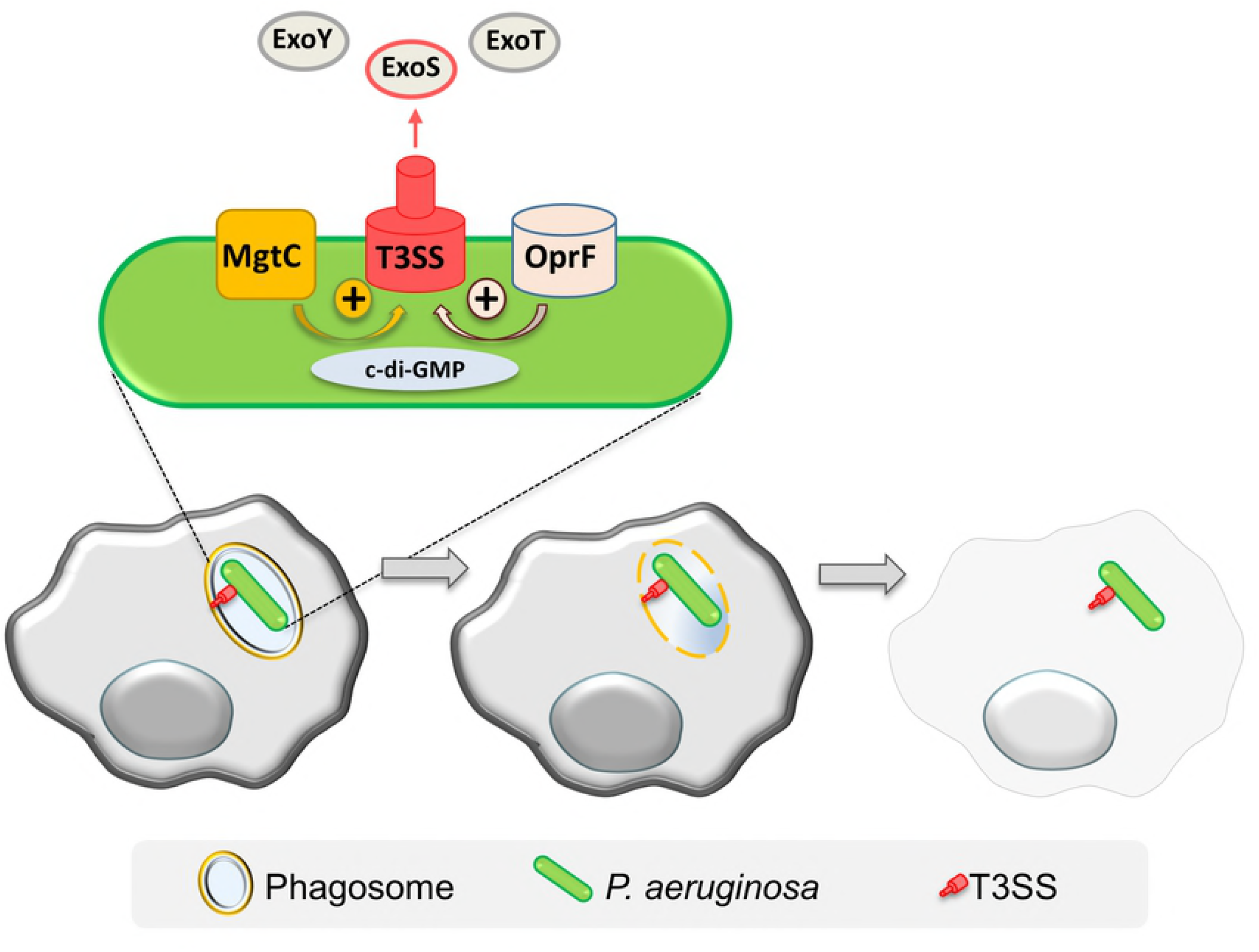
Assessment of access of *P. aeruginosa* to host cytosol using phagosome escape assay. J774 macrophages were infected with PAO1 WT, *ΔmgtC, ΔoprF* and *ΔpscN* strains. After phagocytosis, cells were stained with CCF4-AM in presence of gentamicin. 2 hours post-phagocytosis, the cells were imaged with 10X objective using FITC and DAPI channels. Upon escape of bacteria from phagosome to the cytosol, the CCF4-AM FRET is lost, producing blue color. A. Representative pictures are shown with the following channels: Total cell population is shown in merged green and blue cells, whereas cells with cleaved CCF4 probe are shown in blue in the lower panel. Scale bar is equivalent to 50 μm. B. Images were analyzed and quantified by Cell Profiler software to calculate the percentage of blue cells out of total number of green cells. At least 200 cells were counted per strain. Error bars correspond to standard errors from three independent experiments. The asterisks indicate *P* values (One way ANOVA, where all strains were compared to WT using Dunnet’s multiple comparison post-test, ***P <0.001), showing statistical significance with respect to WT. C. Electron micrograph showing a disrupted vacuole membrane (white arrows) in a macrophage infected with PAO1 strain. The right panel shows higher magnification of the black square in the left panel.

Cumulatively, our results support a T3SS-dependent vacuolar escape for *P. aeruginosa,* leading to the localization of bacteria in the cytoplasm and cell lysis as depicted in the proposed model (Fig 8).

**Fig 8.**
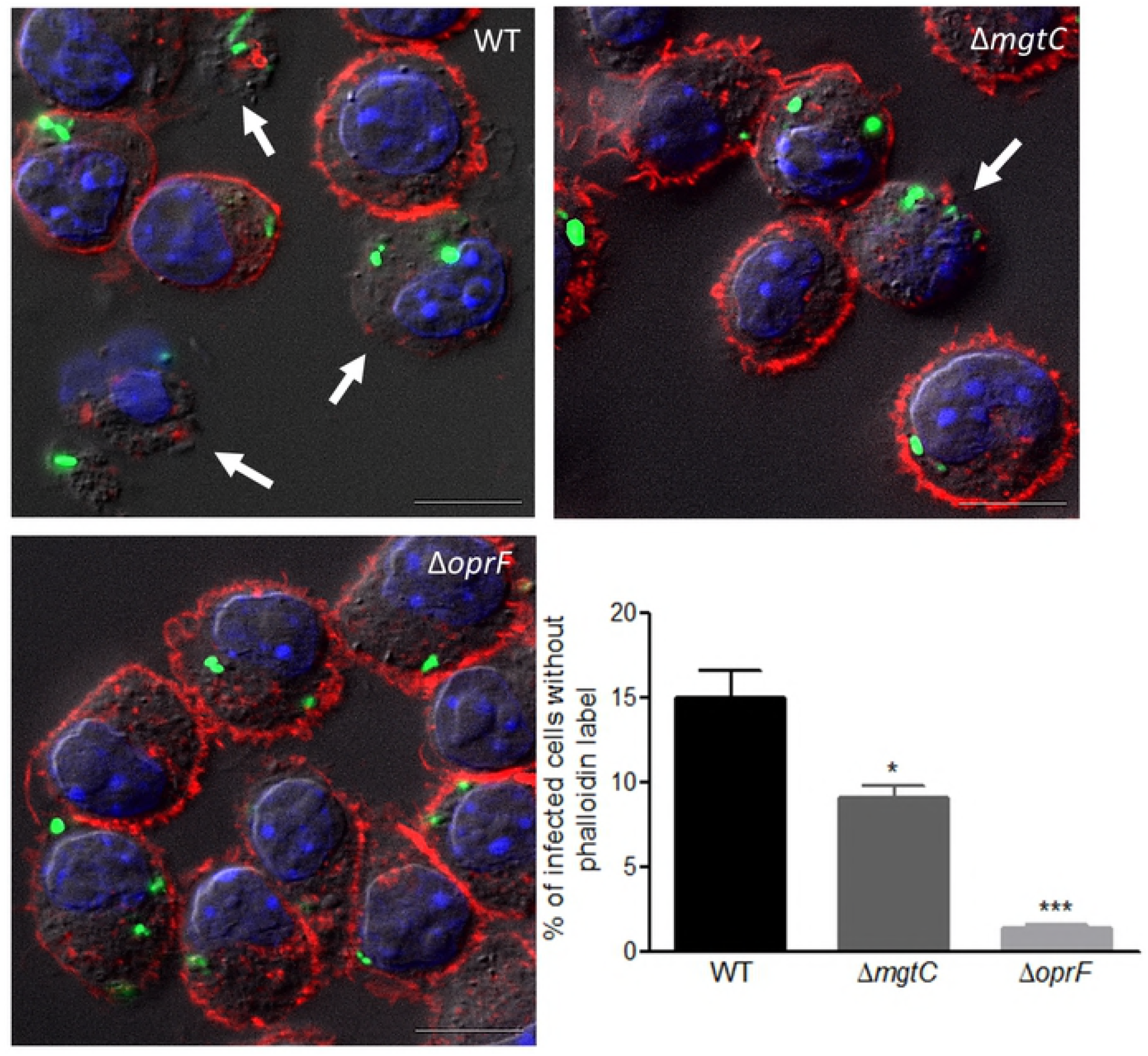
Model for intramacrophage fate of *P. aeruginosa.* Phagocytosed *P. aeruginosa* PAO1 first resides in a vacuole, before escaping the phagosome and promoting macrophage lysis. This cell lysis driven by intracellular *P. aeruginosa* involves the T3SS and more specifically ExoS. MgtC and OprF act positively on the expression of T3SS, possibly by reducing c-di-GMP level, a negative regulator of T3SS expression. Thereby T3SS and its effector ExoS play a role in phagosomal escape and cell lysis.

## Discussion

The ability of professional phagocytes to ingest and kill microorganisms is central to innate immunity and host defense. *P. aeruginosa* is known to avoid being killed by phagocytes through the destruction of immune cells extracellularly as well as avoidance of phagocytosis [19]. However, *P. aeruginosa* has been reported to be engulfed by macrophages in animal infection models [20,21]. In addition, *P. aeruginosa* has been visualized in phagocytes in cell culture models in several studies, where MgtC and OprF, have been shown to be involved in the ability of *P. aeruginosa* to survive in cultured macrophages [23,24]. The virulence of *P. aeruginosa mgtC* mutant can be restored in zebrafish embryos upon macrophages depletion, suggesting that MgtC acts to evade phagocytes [23]. Interestingly, a similar behavior has been reported for a T3SS mutant in the same infection model [20]. We show here that MgtC and OprF regulate T3SS when *P. aeruginosa* resides in macrophages and we describe a novel strategy used by *P. aeruginosa* to escape from macrophages that relies on a T3SS-dependent cell lysis induced by intracellular bacteria (Fig. 8).

Using electron microscopy, we demonstrate that upon phagocytosis, *P. aeruginosa* PAO1 strain resides in membrane bound vacuoles, whereas a cytosolic location can be observed at later time of infection, which corroborates the observation of an otopathogenic *P. aeruginosa* clinical strain in HMDM macrophages [24]. Microscopic analysis of live and fixed cells revealed macrophage lysis driven by intracellular bacteria. This cell lysis is a rapid process, associated with the loss of cortical actin from plasma membrane. We propose that the cell lysis induced by intracellular *P. aeruginosa* is linked to the phagosomal escape of bacteria, which is supported by the observation of cytosolic bacteria and ruptured phagosomal membrane by TEM as well as CCF4-AM/β-lactamase based phagosomal rupture assay.

To better characterize the *P. aeruginosa* factors involved in its intramacrophage fate, we investigated intracellular expression of T3SS genes in *mgtC* and *oprF* mutants, both of which are defective for intramacrophage survival. Expression of T3SS genes upon macrophage infection was significantly decreased in *mgtC* mutant and abrogated in *oprF* mutant. The production of T3SS effectors or needle component was previously known to be altered in the *oprF* mutant [30,32], but a direct effect at the transcriptional level was not investigated before. This regulation could be mediated by c-di-GMP, a known negative regulator of T3SS expression [34], as a reporter assay revealed an increased level of c-di-GMP in both *oprF* and *mgtC* mutants upon monitoring infected macrophages. An increased level of c-di-GMP in *oprF* mutant is consistent with previous results obtained *in vitro* [32]. It is of interest to note that a *Salmonella mgtC* mutant also exhibited increased c-di-GMP level intracellularly [27]. The *P. aeruginosa mgtC* and *oprF* mutants showed a moderate and a more pronounced decrease, respectively, on cytotoxicity driven by intracellular bacteria and phagosomal escape. These phenotypes could be linked to the pattern of T3SS expression in *mgtC* and *oprF* mutants, because a T3SS mutant appeared to lack cytotoxicity driven by intracellular bacteria and showed reduced phagosomal escape.

Our data indicate that the T3SS-mediated cytotoxicity driven by intracellular *P. aeruginosa* is largely dependent on the ExoS effector. In addition, the extent of phagosomal escape of *pscN* and *exoSTY* mutants was similar to that of *exoS* mutant alone, suggesting that ExoS is the main effector protein involved in the exit of *P. aeruginosa* from the phagosome. ExoS, which has GTPase activating protein and ADP ribosyltransferase activities [9], is known for intracellular role in cell types other than macrophages. The T3SS and ExoS are indeed key factors for intracellular *P. aeruginosa* in epithelial cells [17,18]. ExoS and ExoT also promote bacterial survival in neutrophils, by having a protective role in avoiding NADPH-oxidase activity [42]. However, the effect of T3SS towards macrophages was so far restricted to the well-known cytotoxicity caused by extracellular *P. aeruginosa* [37,43–45]. In P. *aeruginosa* strains lacking ExoU toxin, such as PAO1, this cytotoxicity is due to inflammasome activation, which is dependent on T3SS translocation apparatus, but independent of the ExoS effector [37–39,46], thus contrasting with the T3SS mediated cytotoxicity driven by intracellular bacteria described in our study. Hence, the intracellular implication of ExoS suggests that the intracellular life of *P. aeruginosa* in macrophages shares features with the intracellular life of the pathogen in epithelial cells, but with a different outcome. In epithelial cells, ExoS has been involved in avoidance of acidified compartment and intracellular replication in bleb niches [17,18]. Considering our findings, ExoS may allow avoidance of acidification in macrophages and may also contribute to the vacuolar escape in epithelial cells. The intracellular fate of *P. aeruginosa,* however, appears to differ between epithelial cells and macrophages. Whereas bacteria actively replicate in epithelial cells, within bleb niches or in the cytosol [16], replication of bacteria is barely seen within macrophages resulting instead in cell lysis. We propose that upon phagosomal rupture, the cytoplasmic location of *P. aeruginosa* induces inflammasome, possibly through the release of bacterial lipopolysaccharides (LPS) in the cytoplasm, which would promote cell death. Therefore, in addition to the inflammasome activation caused by extracellular *P. aeruginosa* and, as very recently described, by intracellular T3SS-negative *P. aeruginosa* in a context of long-term infection [47], our study suggests that inflammasome and subsequent macrophage death can also be caused by intracellular T3SS-positive *P. aeruginosa.*

In conclusion, our results indicate that *P. aeruginosa* shares common feature with other so-called extracellular pathogens, such as *S. aureus*, which can reside transiently within macrophages [3] and require bacterial factors to survive this stage [48]. A main issue now is to better evaluate the contribution of intramacrophage stage to disease outcome during *P. aeruginosa* infection. Survival of *P. aeruginosa* within macrophages and subsequent bacterial release may play a role in the establishment and dissemination of infection. There is also evidence that intracellular survival may also contribute to persistence of the infection by creating a niche refractory to antibiotic action [22], highlighting the potential importance of this overlooked phase of *P. aeruginosa* infection.

## Materials and methods

### Bacterial strains and growth conditions

Bacterial strains and plasmids are described in Table 1. *P. aeruginosa* mutant strains have been described and phenotypically characterized previously (Table 1). *P. aeruginosa* was grown at 37°C in Luria broth (LB). Plasmid pMF230 expressing GFP constitutively [49] (obtained from Addgene), was introduced in *P. aeruginosa* by conjugation, using an *E. coli* strain containing pRK2013. Recombinant bacteria were selected on *Pseudomonas* isolation agar (PIA) containing carbenicillin at the concentration of 300 μg/ml.

**Table 1.** Bacterial strains and plasmids used in the study

| Name | Description | Reference |
| --- | --- | --- |
| PAO1 |  | Laboratory collection |
| PAO1 $\Delta mgtC$ | | [23] |
| PAO1 $\Delta oprF$ | | [32] |
| PAO1 $\DeltapscN$ | T3SS ATPase mutant | [36] |
| PAO1 $\Delta exoS$ | | [45] |
| PAO1 $\Delta exoSTY$ | | [45] |
| pRK2013 | Tra<sup>+</sup>, Mob<sup>+</sup>, ColE1, Km<sup>r</sup> | Laboratory collection |
| pMF230 | GFP$mut2$, Amp<sup>r</sup> | [49] |
| pTn7CdrA::*gfp*<sup>C</sup> | Amp<sup>r</sup>, Gm<sup>r</sup> | [35] |

### Infection of macrophages and quantification of intracellular bacteria

J774 cells (mouse macrophage cell line J774A.1, gifted by Gisèle Bourg, Inserm U 1047, Nîmes, France) were maintained at 37°C in 5% CO_2_ in Dulbecco’s modified Eagle medium (DMEM) (Gibco) supplemented with 10% fetal bovine serum (FBS) (Gibco). The infection of J774 macrophages by *P. aeruginosa* was carried out essentially as described previously [23]. Mid-log phase *P. aeruginosa* grown in LB broth was centrifuged and resuspended in PBS to infect J774 macrophages (5×10^5^ cells/well) at an MOI of 10. After centrifugation of the 24-well culture plate for 5 min for synchronization of infection, bacterial phagocytosis was allowed for 25 min. Cells were washed three times with sterile PBS and fresh DMEM medium supplemented with 400 μg/ml gentamicin was added and retained throughout the infection. To quantify internalized bacteria, macrophages were lysed after 20 min (T0) or 2 hrs (T1) of gentamicin treatment, by using 0.1% Triton X-100 and the number of viable bacteria was determined by subsequent plating onto LB agar plates. The percentage of survival was obtained by multiplying with 100 the ratio of the bacterial CFU at T1 to that of T0.

### Live microscopy

J774 macrophages were seeded in ibidi μ slide (8 wells) in DMEM medium supplemented with 10% FBS and infected with *P. aeruginosa* PAO1 expressing GFP as described in the previous section. Imaging started after 30 min of phagocytosis, when the media was changed to DMEM supplemented with gentamicin until 3 hrs post phagocytosis. Cells were imaged using an inverted epifluorescence microscope (Axioobserver, Zeiss), equipped with an incubation chamber set-up at 37°C and 5% CO_2_ and a CoolSNAP HQ2 CCD camera (Photometrics). Time-lapse experiments were performed, by automatic acquisition of random fields using a 63X Apochromat objective (NA 1.4). The frequency of acquisition is indicated in figure legends. Image treatment and analysis were performed using Zen software (Zeiss).

### Transmission Electron microscopy

Macrophages were seeded on glass coverslips and infected as described above. Infected cells were fixed for 4 hrs at room temperature with 2.5% gluteraldehyde in cacodylate buffer 0.1 M pH 7.4 with 5mM CaCl_2_, washed with cacodylate buffer, post-fixed for 1 hr in 1% osmium tetroxide and 1.5 % potassium ferricyanid in cacodylate buffer, washed with distilled water, followed by overnight incubation in 2% uranyl acetate, prepared in water. Dehydration was performed through acetonitrile series and samples were impregnated in epon 118: acetonitrile 50:50, followed by two times for 1 hr in 100% epon. After overnight polymerization at 60°C, coverslips were detached by thermal shock with liquid nitrogen. Polymerization was then prolonged for 48 hrs at 60°C. Ultrathin sections of 70 nm were cut with a Leica UC7 ultramicrotome (Leica microsystems), counterstained with lead citrate and observed in a Jeol 1200 EXII transmission electron microscope. All chemicals were from Electron Microscopy Sciences (USA) and solvents were from Sigma. Images were processed using Fiji software.

### Colocalization of *P. aeruginosa* with acidic compartments

Macrophages were infected with GFP labelled *P. aeruginosa* as described above. After 2.5 hrs of gentamicin treatment, infected J774 cells were washed twice with PBS and incubated with 50 nM Lysotracker red DND-99 (Molecular Probes) in DMEM for 10 min to stain lysosomes. Cells were then washed with PBS and fixed with 4% paraformaldehyde in PBS and mounted on glass slides in Vectashield with DAPI (Vector Laboratories, Inc). The slides were examined using an upright fluorescence microscope (Axioimager Z2, Zeiss) equipped with an Apotome 1 for optical sectioning. A 63X Apochromat Objective (NA 1.4) was used, transmitted light was acquired using differential interference contrast (DIC), FITC filter was used to visualize GFP expressing bacteria and Lysotracker red fluorescence was acquired using a texas red filter set.

### Phalloidin labeling

J774 macrophages were seeded on glass coverslips and infected with GFP expressing bacteria as described in the previous sections. For cytochalasin treatment, DMEM containing 2 μM cytochalasin D (Sigma) was added to the macrophages 1 hour before infection and maintained during phagocytosis. After phagocytosis, the cells were maintained in 1 μM cytochalasin D till the end of the experiment. For untreated control, 0.2 % DMSO (solvent control) in DMEM was added to the cells before and during phagocytosis. After phagocytosis, cells were maintained in 0.1 % DMSO. After fixation with 4% paraformaldehyde (EMS, USA) in PBS for 5 min, cells were washed once with PBS and permeabilized by adding 0.1% triton X-100 for 1 min 30 sec. Cells were then washed once with PBS and incubated with 1 μg/ml Tetramethylrhodamine B isothiocyanate (TRITC)-labeled phalloidin (Sigma-Aldrich) in PBS for 30 min in dark. Cells were washed twice with PBS and coverslips were mounted on glass slides in Vectashield with DAPI (Vector Laboratories, Inc). The slides were examined using an upright fluorescence microscope (Axioimager Z1, Zeiss) equipped with an Apotome 1 for optical sectioning. A 63X Apochromat Objective (NA 1.4) was used, transmitted light was acquired using DIC. FITC and texas red filters were used to visualize GFP expressing bacteria and phalloidin respectively. Images were processed using ZEN software (Zeiss). Cells were counted manually, where infected cells lacking the phalloidin stain were considered as lysed. Percentage of such lysed cells with intracellular bacteria out of total number of infected cells was calculated and plotted for each strain.

### LDH cytotoxicity assay

The cytotoxicity was assessed by release of lactate dehydrogenase (LDH) from infected J774 macrophages infected, using the Pierce LDH cytotoxicity assay kit (Thermo Scientific). Macrophages were infected for 2 hrs at an MOI of 10 as described above, except that cells were seeded in a 96 well plate and extracellular bacteria were not removed. The assay was performed on 50 μl of the culture supernatant according to manufacturer’s instructions. LDH release was obtained by subtracting the 680 nm absorbance value from 490 nm absorbance. The percentage of LDH release was first normalized to that of the uninfected control and then calculated relatively to that of uninfected cells lysed with Triton X-100, which was set at 100% LDH release.

### RNA extraction and quantitative RT-PCR (qRT-PCR)

For bacterial RNA extraction from infected J774, 6.5×10^6^ macrophages were seeded into a 100 cm^2^ tissue culture dish and infected at an MOI of 10 as described above. 1 hour after phagocytosis, cells were washed three times with PBS, lysed with 0.1% Triton X100 and pelleted by centrifugation at 13000 rpm for 10 min at 15°C. Bacteria were resuspended in 500 μl PBS and the non resuspended cellular debris was discarded. 900 μl of RNA protect reagent (Qiagen) was added and incubated for 5 min. The sample was centrifuged at 13000 rpm for 10 min. Bacteria in the pellet were lysed with lyzozyme and RNA was prepared with RNEasy kit (Qiagen). Superscript III reverse transcriptase (Invitrogen) was used for reverse transcription. Controls without reverse transcriptase were done on each RNA sample to rule out possible DNA contamination. Quantitative real-time PCR (q-RT-PCR) was performed using a Light Cycler 480 SYBR Green I Master mix in a 480 Light Cycler instrument (Roche). PCR conditions were as follows: 3 min denaturation at 98°C, 45 cycles of 98°C for 5 sec, 60°C for 10 sec and 72°C for 10 sec. The sequences of primers used for RT-PCR are listed in Table S1.

### Cyclic di-GMP reporter assay

The strains were transformed by electroporation with the plasmid pCdrA::*gfp* [32,35], which expresses GFP under the control of the promoter of *cdrA,* a c-di-GMP responsive gene, and carries a gentamicin resistance gene. Overnight cultures, grown in LB with 100 μg/ml gentamicin, were subcultured in LB. These cultures were used to infect J774 cells seeded in a 96 well plate (Greiner, Flat-Bottom), containing 10^5^ cells per well with an MOI of 20 after normalizing the inoculum to their OD_600_. After phagocytosis, DMEM containing 300 μg/ml of amikacin, instead of gentamicin, was added to eliminate extracellular bacteria, as these strains are resistant to gentamicin. At the required time point, Tecan fluorimeter (Spark 20M) was used to measure fluorescence (excitation, 485 nm and emission, 520 nm) of cells at the Z point where emission peak could be obtained in comparison to the blank. Fluorescence was plotted for each strain in terms of arbitrary units (AU).

### CCF4 fluorometric assay to monitor the escape of *P. aeruginosa* in host cytosol

The vacuole escape assay was adapted from the CCF4 FRET assay [40] using the CCF4-AM LiveBlazer Loading Kit (Invitrogen). Briefly, J774 macrophages were seeded in 96 well plate (Greiner, Flat-Bottom), containing 5×10^4^ cells per well. Overnight bacterial cultures were subcultured in LB with 50 μg/ml of ampicillin to enhance the expression of beta-lactamase, present naturally in *P. aeruginosa*. Infection was carried out as mentioned in the previous sections at the MOI of 10. After phagocytosis, the cells were washed thrice with PBS to remove extracellular bacteria. 100 μl of HBSS buffer containing 3 mM probenecid and gentamicin (400 μg/ml), was added in each well. The substrate solution was prepared by mixing 6 μl of CCF4-AM (solution A), 60 μl of solution B and 934 μl of solution C. 20 μl of the substrate solution was added to each well and the plate was incubated in dark at 37 °C with 5% CO_2_. After 2 hrs, the cells were imaged as described in the Live Imaging section, using a 10X objective. GFP and DAPI channels were used to visualize CCF4-FRET (Green) and loss of FRET (Blue) respectively. Each sample was taken in triplicate and image acquisition was performed by automated random acquisition. Images were analyzed by Cell Profiler software to calculate the number of blue cells out of green cells. The percentage of blue cells, representing the cells with cytosolic bacteria, out of total green cells was plotted. All strains were tested for their ability to cleave CCF4 *in vitro*, before carrying out the vacuole rupture assay. Overnight cultures grown in LB were subcultured at the ratio of 1:20 in LB with 100 μg/ml of ampicillin. After 2 hours of growth the cultures were centrifuged and resuspended in PBS. 100 μl of this was aliquoted in 96 well plate (Greiner, Flat-Bottom) and 20 μl of CCF4 subtrate solution (A+B+C) was added. Tecan fluorimeter (Spark 20M) was used to measure fluorescence (excitation, 405 nm and emission, 535 nm for green and 450 nm for blue) using PBS as blank. Blue fluorescence was observed for all strains and none of the mutants exhibited lower value than the wild-type strain.

## Acknowledgments

We thank Sylvie Chevalier (Rouen) and Ina Attrée (Grenoble) for providing strains and plasmids, the Montpellier RIO Imaging microscopy platform for photonic Microscopy and the Electron Microscopy facility of the University of Montpellier (MEA) for sample preparation and transmission electron microscopy. We thank Bérengère Ize and Laila Gannoun-Zaki for critical reading of the manuscript, Matteo Bonazzi and Fernande Siadous for their expertise with the CCF4 assay.

## Supporting Information

**S1 Fig. Live imaging of macrophages infected with *P. aeruginosa*.** J774 macrophages were infected with PAO1 wild-type strain expressing GFP. Time lapse imaging was started at 1.5 hrs post-phagocytosis. Cells were maintained in DMEM supplemented with gentamicin, at 37 °C and 5% CO_2_ throughout imaging. White arrows point at infected cells that undergo lysis, whereas black arrows indicate uninfected cells that do not lyse. Images were taken between 1.5 hrs and 3 hrs post-phagocytosis as shown on the panels. Scale bar is equivalent to 10 μm.

**S2 Fig. Colocalization of *P. aeruginosa* with a probe that labels acidic compartments.** J774 macrophages were infected with PAO1 expressing GFP. After 2.5 hrs of gentamicin treatment, infected J774 cells were incubated with Lysotracker for 10 min, a red fluorescent weak base that accumulates in acidic compartments. Cells were then fixed and imaged with fluorescence microscope. (A) The image shows individual panels for Differential Interference Contrast (DIC), lysosomal compartment (red), bacteria expressing GFP (green), the nucleus (blue) and merged image of all channels. The solid arrow shows colocalization of bacteria with lysotracker and dashed arrow shows non-colocalization. Scale bar is equivalent to 5 μm. (B) 3D-reconstructed image of the same area of (A).

**S3 Fig. Survival of bacteria upon phagocytosis.** J774 macrophages were infected with PAO1 WT, *ΔmgtC* and *ΔoprF* strains. After phagocytosis, cells were maintained in DMEM supplemented with gentamicin. 2 hours post-phagocytosis, infected cells were lysed and intracellular bacteria were plated on LB agar plates to obtain CFU values. Results are expressed as percentage of surviving bacteria at the time T1 (2 hrs after phagocytosis) compared to the number of bacteria at time T0 (20 min after phagocytosis). Error bars correspond to standard errors (SE) from four independent experiments. The asterisks indicate *P* values (One way ANOVA, where all strains were compared to WT using Dunnet’s multiple comparison post-test, **P <0.01), showing statistical significance with respect to WT.

**S4 Fig. Visualization (A) and quantification (B) of lysed infected cells upon cytochalasin D treatment.** GFP expressing PAO1 used for infecting J774 macrophages. DMEM containing 2 μM cytochalasin D (Sigma) was added to the macrophages 1 hour before infection and maintained during phagocytosis. After phagocytosis, cells were maintained in 1 μM cytochalasin in DMEM supplemented with gentamicin till the end of the experiment. For untreated control, 0.2 % DMSO (solvent control) in DMEM was added to the cells before and during phagocytosis. After phagocytosis, cells were maintained in 0.1 % DMSO in DMEM with gentamicin, fixed 2 hrs post-phagocytosis, stained with phalloidin and imaged with fluorescent microscope. DAPI was used to stain the nucleus. Cells that have intracellular bacteria, but lack the phalloidin cortical label were considered as lysed by intracellular bacteria (shown by arrows). Scale bar is equivalent to 10 μm. Percentage of lysed cells with intracellular bacteria out of total number of cells was plotted. Error bars correspond to standard errors from two independent experiments. At least 200 cells were counted per strain. The asterisks indicate *P* values (Student’s t-test, **P <0.01), showing statistical significance with respect to DMSO control.

**S5 Fig. Quantification of cell lysis driven by extracellular bacteria.** Release of LDH was measured from J774 macrophages infected for 2 hrs with PAO1 WT, *ΔpscN, ΔexoS* and *ΔexoSTY* mutant strains to quantify the cytotoxicity. The percentage of LDH release was calculated relatively to that of total uninfected cells lysed with Triton X-100, which was set at 100% LDH release. Error bars correspond to standard errors (SE) from at least four independent experiments. The asterisks indicate *P* values (One way ANOVA, where all strains were compared to WT using Dunnet’s multiple comparison post-test, **P <0.01), showing statistical significance with respect to WT.

**S6 Fig. Phagosome escape assay of T3SS mutants.** J774 macrophages were infected with PAO1 WT, *ΔpscN, ΔexoS* and *ΔexoSTY* strains. After phagocytosis, cells were stained with CCF4-AM in presence of gentamicin. 2 hours post-phagocytosis, the cells were imaged with 10X objective using FITC and DAPI channels. Upon escape of bacteria from phagosome to the cytosol, the CCF4-AM FRET is lost, producing blue color. Images were analyzed and quantified by Cell Profiler software to calculate the percentage of blue cells out of total number of green cells. At least 200 cells were counted per strain. Error bars correspond to standard errors from four independent experiments. The asterisks indicate *P* values (One way ANOVA, where all strains were compared to WT using Dunnet’s multiple comparison posttest, *P <0.05), showing statistical significance with respect to WT.

**S1 Movie. Live microscopy movie imaging cell lysis in real time.** J774 macrophages were infected with PAO1 WT strain expressing GFP. Cells were maintained in DMEM supplemented with gentamicin, at 37 °C and 5% CO_2_ throughout imaging. Imaging was started at 3 hrs post-phagocytosis and continued for 10 min with interval of 30 sec between frames. The time frame is displayed in the movie in the format of mm:ss. The movie shows quick lysis of macrophages harboring intracellular bacteria occurring within a minute.

**S1 Table. List of primers used for RT-PCR.**

